# Predicting the activation of the androgen receptor by mixtures of ligands using Generalized Concentration Addition

**DOI:** 10.1101/2020.05.02.074112

**Authors:** Jennifer J Schlezinger, Wendy Heiger-Bernays, Thomas F Webster

## Abstract

Concentration/dose addition (CA) is widely used for compounds that act by similar mechanisms. But, CA cannot make predictions for mixtures of full and partial agonists for effect levels above that of the least efficacious component. As partial agonists are common, we developed Generalized Concentration Addition (GCA), which has been successfully applied to systems in which ligands compete for a single binding site. Here, we applied a pharmacodynamic model for a system with two binding sites, the androgen receptor (AR). AR acts according to the classic homodimer activation model: each cytoplasmic AR protein binds ligand, undergoes a conformational change that relieves inhibition of dimerization, and binds to DNA response elements as a dimer. We generated individual dose-response data for full (dihydroxytestosterone, BMS564929) and partial (TFM-4AS-1) agonists and a competitive antagonist (MDV3100) using reporter data generated in the MDA-kb2 cell line. We used the Schild method to estimate the binding affinity of AR for MDV3100. Data for individual compounds fit the AR pharmacodynamic model well. The partial agonist had agonistic effects at low effect levels and antagonistic effects at high levels, as predicted by pharmacological theory. The GCA model fit the empirical mixtures data—full/full agonist, full/partial agonist and full agonist/antagonist—as well or better than relative potency factors (a special case of CA) or effect summation. The ability of generalized concentration addition to predict the activity of mixtures of different types of androgen receptor ligands is important as a number of environmental compounds act as partial AR agonists or antagonists.

## Introduction

Nuclear receptors (NRs) are ligand-activated transcription factors and are the mechanism by which lipophilic hormones and hormone-like molecules regulate gene expression. Thus, they are critical targets of endocrine disrupting chemicals (EDCs). Androgen receptor (AR) is an NR that binds ligands including hormones (e.g., testosterone), pharmaceuticals (e.g., flutamide) and environmental EDCs (e.g., cyprodinil, vinclozolin, procymidone). In females, AR signaling is necessary for efficient folliculogenesis (i.e., AR knockout females have impaired folliculogenesis and display early onset infertility) (Matsumoto *et al.*, 2008). In males AR signaling is necessary for the reproductive function (i.e., development of reproductive organs, spermatogenesis, and brain masculinization) and regulates bone mass, muscle mass, fat distribution and hair patterning (Matsumoto, Shiina, Kawano, Sato and Kato, 2008). There is evidence for a decline in human sperm counts between 1973 and 2011 (Barratt *et al.*, 2018), and an increased incidence of symptoms of testicular dysgenesis syndrome, including testicular cancer, disorders of sexual development, cryptorchidism, hypospadias (Skakkebaek *et al.*, 2016). Given the importance of AR signaling in physiology, increasing human exposure to environmental chemicals that are AR ligands and declining human reproductive health, it is necessary to be able to model how multiple ligands interact to activate AR.

An approach commonly used to assess the effect of mixtures of EDCs is Concentration Addition, also called dose addition (CA)(Rajapakse *et al.*, 2004). This model requires that all components of the mixture have the same efficacy or maximal response. However, ligands of NRs may act as full and partial agonists or competitive antagonists. Partial agonists are ligands that produce a lower maximal response than full agonists, and antagonists do not induce a response. We developed GCA to resolve the inability of CA to make predictions for mixtures including partial agonists and/or antagonists. GCA uses inverse mathematical functions allowing for some components to have submaximal efficacy (Howard and Webster, 2009). GCA successfully predicts such mixture effects for ligand-receptor systems with one binding site. (e.g., AhR, PPARγ) (Howard *et al.*, 2010; Watt *et al.*, 2016). We recently developed a mathematical dose response model for receptors that bind ligand at each monomer and then dimerize and showed that it fulfilled the requirements for GCA (Webster and Schlezinger, 2019).

Here, we test the ability of GCA to model the activation of AR by ligand mixtures and compare it to effect summation (ES), a classic additivity model, as well as relative potency factors (RPF), a special case of CA and GCA. Using MDA-kB2 cells, which express a steroid response element-dependent luciferase reporter (Wilson *et al.*, 2004), we generated dose-responses for individual synthetic AR ligands. We predicted mixture effects using the different additivity models and compared these predictions with empirical results. We show that the dissociation constant derived from the Schild method can be incorporated into GCA in order to model mixtures containing antagonists. We establish GCA as a model for predicting AR activation by mixtures containing diverse AR ligand types.

## Materials and Methods

### Chemicals

BMS564929 (CAS # 627530-84-1, Cat. # 5274, purity ≥ 98%) and TFM-4AS-1 (CAS # 188589-61-9, Cat. # 3813, purity ≥ 98%) were purchased from Tocris Bioscience (Bristol, UK). MDV3100 (CAS # 915087-33-1, Cat. # sc-364354, purity ≥ 98%) was purchased from Santa Cruz Biotechnology (Dallas, TX). Dihydroxytestosterone (CAS # 521-18-6, Cat. # 15874, purity ≥ 97%) was purchased from Cayman Chemical (Ann Arbor, MI). All other reagents were purchased from Thermo Fisher Scientific (Waltham, MA), unless indicated.

### AR Reporter Assays

MDA-kb2 cells (RRID:CVCL_6421, CRL-2713; ATCC, Manassas, VA) were maintained in Leibovitz’s L-15 medium (ATCC, Cat. # 30-2008) supplemented with 10% fetal bovine serum (Gemini Bio-Products, West Sacramento, CA) and Antibiotic-Antimycotic (100 U/ml penicillin, 100 µg/ml streptomycin, 0.25 µg/ml amphotericin B) at 37°C without CO_2_ supplementation. Seven days prior to plating for an experiment, the cultures were switched to steroid-free medium (Leibovitz’s L-15 medium supplemented with 10% charcoal/dextran stripped fetal bovine serum and Antibiotic-Antimycotic). The medium was replaced once prior to experiment plating.

For experiments, MDA-kb2 cells were plated at 40,000 cells per well in 0.2 ml steroid-free medium in white-sided, 96-well plates. Following a 48 hr incubation, cells received no treatment (medium only), Vh (DMSO, 0.25% for individual dose responses or 0.5% for combination dose responses), the positive control (dihydroxytestosterone (DHT), 1 × 10^−8^ M) or AR ligands at a range of concentrations either alone or in combination (DHT, 1 × 10^−13^ M – 1 × 10^−6^ M; BMS564929 1 × 10^−12^ M – 1 × 10^−6^ M; TFM-4AS-1 1 × 10^−11^ M – 4 × 10^−4^ M; MDV3100 1 × 10^−10^ M – 2 × 10^−5^ M). Vh and experimental chemicals were applied to duplicate wells. The positive control was applied to six wells of every experimental plate. Six wells were left untreated (Naïve) in every experimental plate. After 24h of treatment, cells were analyzed using the Steady-Luc™ Firefly HTS Assay Kit (Biotium, Freemont, CA). Luminescence was measured using a Synergy2 plate reader (Biotek Inc, Winooski, VT). To assess toxicity, MTS reagent was added 4 hours prior to analysis of absorbance at 490 nm (cat. #ab197010, Abcam, Cambridge, UK). Experiments were repeated at least 4 times.

Data from replicate wells were averaged prior to analysis. Data were normalized using a method to minimize intra- and inter-experimental variation (Rajapakse, Silva, Scholze and Kortenkamp, 2004), as follows:

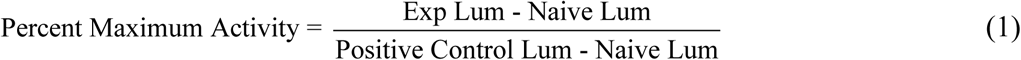

AR activation is reported as “% Maximum Activity.”

### Schild Method

The Schild method was carried out as described (Kenakin, 2014). Using the MDA-kb2 experimental approach described above, dose response data for the full agonist BMS564929 (1 × 10^−11^ M - 1 × 10^−7^ M) were generated in the presence of Vh or the competitive antagonist MDV3100 (4 × 10^−8^, 1 × 10^−7^, 4 × 10^−7^, 1× 10^−6^, 2 × 10^−6^ or 4 × 10^−6^ M). Data from 6 experiments were averaged. The means for each dose response curve were fit with the “log(agonist) vs response – Variable Slope (four parameters)” Hill function in Prism (version 6, GraphPad, San Diego, CA); maximum, slope and sum of squares values were determined from each fit. The mean of the maxima and of the slopes of each of the 7 dose response curves were calculated. The dose response curves then were refitted using the “log(agonist) vs response – Variable Slope (four parameters)” function but were constrained by the means of the maxima and the slopes, and the sum of squares was determined.

The fits of the unconstrained and constrained models then were examined using Akaike’s information criterion for small sample sizes (AICc)(Portet, 2020), where:

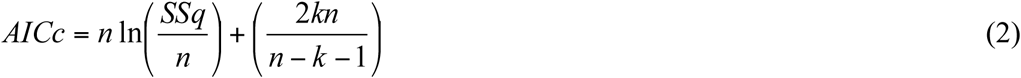

where *n* is the sample size, *SSq* in the sum of squared deviations between the model and the actual responses, and *k* is the number of parameters.

Using the data from the model (constrained) with the lower AIC, we calculated the dose ratio (DR) for each dose response curve by dividing the EC_50_ of each curve by the EC_50_ of the curve generated in the absence of MDV3100. The Schild plot was generated by plotting the log (DR-1) versus the MDV3100 concentration. We assessed the suitability of the Schild plot by ensuring that the slope was greater than 0.8, that there were data near the log (DR-1) = 0, and that a wide concentration range was represented (Kenakin, 2014).

### Hill and Pharmacodynamic Models

Dose response data for each individual compound except the competitive antagonist were modeled with a four parameter Hill function (equation 3)

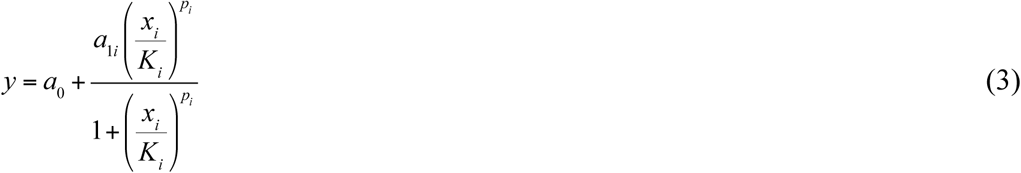

where *y* is the response and *x*_*i*_ is the concentration of ligand *i*, and the four parameters are as follows: the background response in the absence of ligand (*a*_0_,); the span, the maximum response above background (*a*_1i_); the EC_50_ (*K*_*i*_), and the slope parameter (*p*_*i*_). The latter three may differ by compound.

Derivation of the pharmacodynamic model (PDM) for homodimers such as the androgen receptor was described previously (Webster and Schlezinger 2019). Briefly, we assume that receptor monomers reversibly bind ligand and dimerize leading to signal. We also assume equilibrium and constant total receptor concentration. The result can be written as a composite function

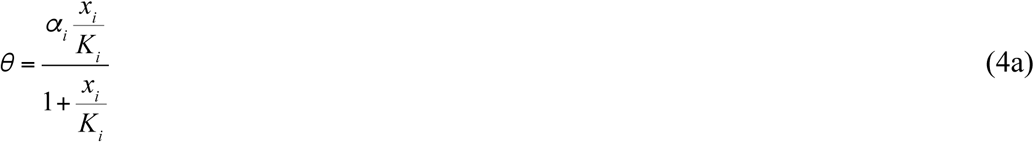

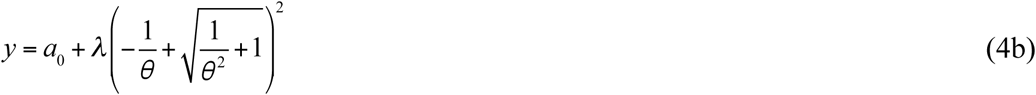

where equation (4a) is a Hill function of slope one, but the compound-specific parameters (*α*_i_, *K*_i_) are interpreted somewhat differently. The second function contains the background response (*a*_0_) and a scaling parameter (*λ*) that does not depend on the compound.

### Generalized Concentration Addition (GCA) for Homodimers

GCA is an extension of concentration addition defined by the following equation

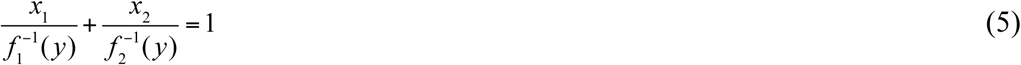

for a binary mixture, easily generalized to more components (Howard and Webster 2009). The denominators are the inverse dose-response functions for the individual compounds. To predict the combined effect of mixtures of compounds differing in efficacy, the inverse function must yield real numbers for partial agonists at effect levels above their maximal effect. As previously shown, this requirement is met by the homodimer PDM (equation 4) but not generally by Hill functions with slope parameters different from one and many other function used for dose-response analysis (Webster and Schlezinger 2019).

Application of GCA replaces (4a) by the following equation for a binary mixture:

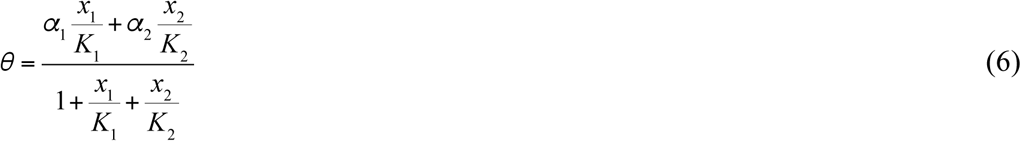

while the outer part of the composite function (4b) remains the same (Webster and Schlezinger 2019). This result is easily generalized to more complicated mixtures. For application to a mixture of an agonist and a competitive antagonist, the alpha parameter for the latter is set to zero.

### Effect Summation (ES)

ES is mixtures model that sums the individual responses (rather than the concentrations) of the mixture components. The prediction of ES for a binary mixture is

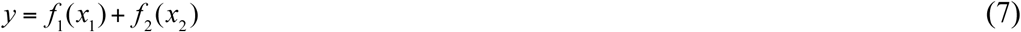

where *f*_i_(*x*_i_) is the dose response for compound *i* alone. Note that effect summation is generally not considered to be a useful model in mixtures toxicology (Berenbaum 1989).

### Relative Potency Factor (RPF)

The relative potency factor model for mixtures assumes that mixture components differ only in their potency. For Hill functions this means that the maximal effects and slope parameters are the same. The relative potency factor is often based on the ratio of the EC50 of a compound compared to a reference compound. The toxic equivalence factors used for dioxin-like compounds are a well known example (Van den Berg et al 2006). The RPF model for a binary mixture is described by

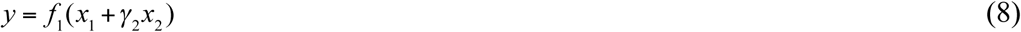

where f_1_(·) is the dose response function for the reference compound (*x*_1_) and γ_2_ is the RPF for compound 2, typically estimated as the EC50 for compound 1 divided by the EC50 for compound 2. The RPF model is a special case of CA, which is in turn a special case of GCA (Howard and Webster 2009). Unlike CA, the RPF model can make predictions for mixtures of full and partial agonists at effect levels above the maximal effect of the partial agonist, but this violates the assumptions underlying the model.

### Statistics and Software

Hill functions were fit to individual compounds using R (R Core Team 2017) as well as using Prism (version 6, GraphPad), yielding similar results (not shown). After subtracting background response, the homodimer PDM model for each compound was fit using R for a range of values of λ yielding estimates of *α*_i_ and *K*_i_. As λ is common across compounds (equation 4b), the value of λ was chosen that minimizes the mean squared error. Dose-response surfaces for mixtures data were generated using R. Predictions for the three mixtures models were made using the estimated parameters. For GCA, the estimated values of *a*_0_, *α*_i_, *K*_i_ and λ were substituted into equations 6 and 4b. Effect summation estimates (equation 7) and relative potency factors (equation 8) were based on the fits of the four parameter Hill function. The dose-response surfaces for each model were compared with empirical data using root mean squared error (RMSE).

## Results

We began by establishing the dose-responses of three AR ligands, DHT (full agonist), BMS564929 (full agonist) and TFM-4AS-1 (partial agonist). MDA-kb2 cells express 240 fmol AR/mg-protein and are stably transfected with a luciferase construct driven by a steroid response element (Hall *et al.*, 1990; Ham *et al.*, 1988; Wilson, Bobseine and Gray, 2004). The assay is included in Tox21 and ToxCast (Kleinstreuer *et al.*, 2017). No toxicity was evident at the concentrations tested (Supplemental Material, Figure S1). Fitted parameters and RMSEs are presented in Tables S1 and S2, respectively. As shown in Figure 1, the dose response data for BMS564929 and TFM-4AS-1 were very well fit by both the four parameter Hill function and the homodimer PDM; the Hill function fit slightly better for DHT.

**Figure 1.**
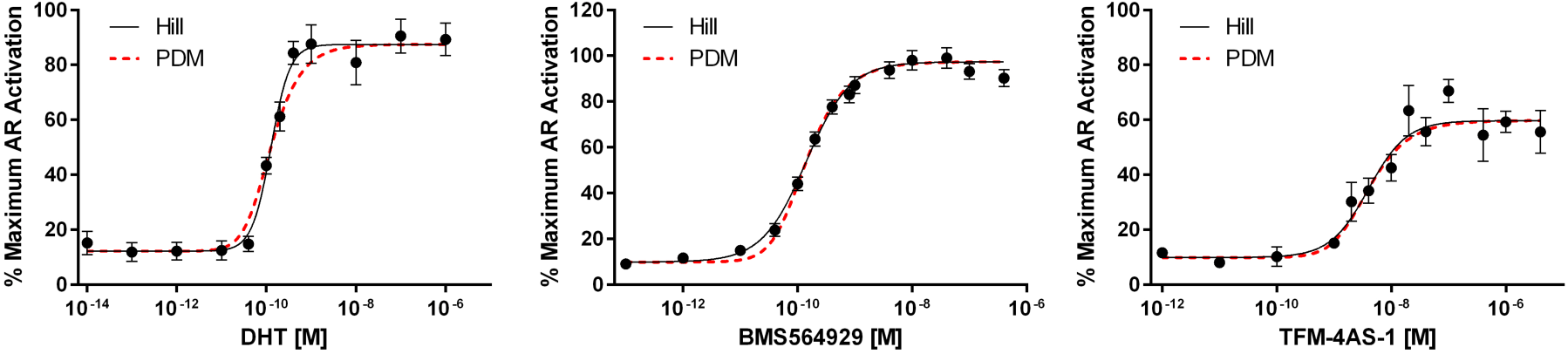
Dose response curves for androgen receptor activation by full and partial agonists. MDA-kb2 cells were plated in steroid-free medium and dosed with Vh (0.5% DMSO), DHT (1 × 10^−13^ M – 1 × 10^−6^ M), BMS564929 (1 × 10^−12^ M – 1 × 10^−6^ M) or TFM-4AS-1 (1 × 10^−11^M – 4 × 10^−4^ M). Luminescence was measured after 24 hrs. Ligand-induced luminescence was normalized to the positive control-induced luminescence. Data from Vh-treated wells are reported as the lowest concentration in each plot. Dose response data were fit with a Hill function or a PDM function. **(A)** Dose response of full agonist DHT. **(B)** Dose response of full agonist BMS564929. **(C)** Dose response of partial agonist TFM-4AS-1. Data are presented as means ± SE of at least 4 independent experiments.

We hypothesized that mixtures of BMS564929 with DHT and with TFM-4AS-1 would show disparate effects on the overall efficacy of AR activation. For mixtures of BMS564929 with DHT, we expected that the efficacy of the mixture would reach 100% because each is a full AR agonist. This is evident, as shown in Figure 2a. For mixtures of BMS564929 with TFM-4AS-1, we expected that the mixture would cause a reduction in overall AR activation at effect levels above the efficacy of TFM-4AS-1, owing to the behavior of a partial agonist to act in the manner of a competitive antagonist in a mixture. Figure 2b shows that at high effect levels, the response to BMS564929 is reduced and the overall activation of AR by the mixture decreases.

**Figure 2.**
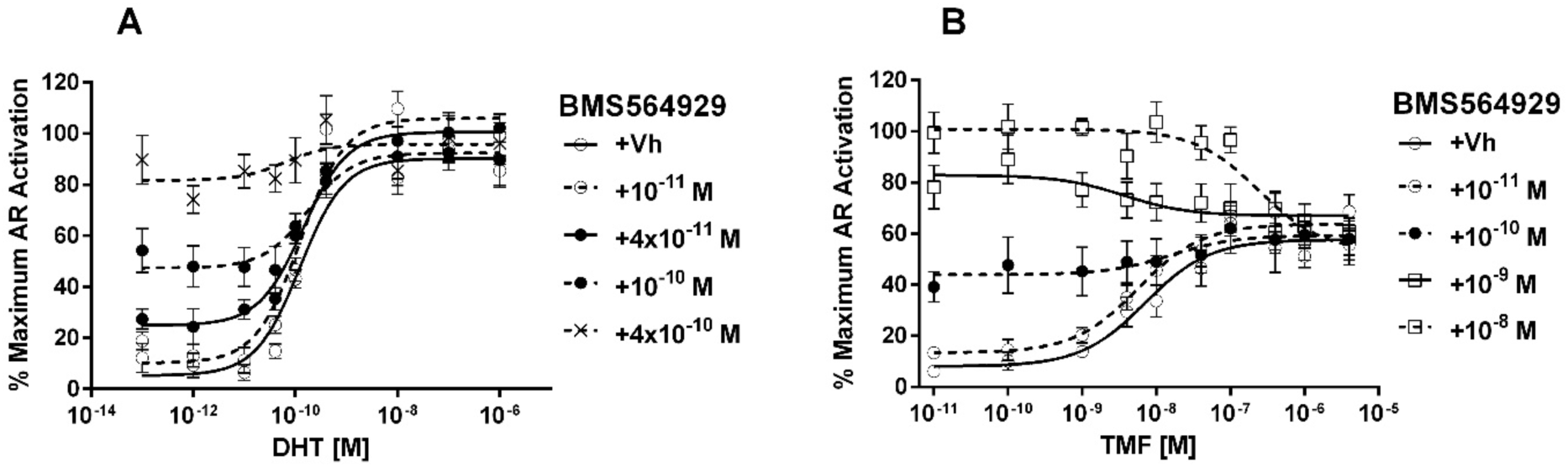
Efficacy of androgen receptor activation by a mixture of a full agonist with a full agonist (A) and with a partial agonist (B). MDA-kb2 cells were plated in steroid-free medium and dosed with Vh (0.5% DMSO, shown as lowest concentration) or BMS564929 (1 × 10^−11^ M – 1 × 10^−8^ M) with DHT (1 × 10^−12^ M – 1 × 10^−6^ M), or with TFM-4AS-1 (1 × 10^−10^ M – 4 × 10^−5^ M). Luminescence was measured after 24 hrs. Ligand-induced luminescence was normalized to the positive control-induced luminescence. Data from Vh-treated wells are reported as the lowest concentration in each plot. Data are presented as means ± SE of at least 4 independent experiments.

To compare the empirical dose response data with prediction of the different mixtures models, three-dimensional response surfaces of AR activation by the full/full and full/partial agonist combinations were plotted. Figure 3 shows that for the full agonist/full agonist mixture, the GCA model fits the experimental data the best, i.e., it has the lowest RMSE, and the RPF model fit almost as well. The latter is expected since RPF is a special case of GCA where compounds differ only in potency, as would be expected for full agonists. The small difference is due to the slightly different maximal effects and slopes for DHT and BMS as shown in Figure 1. As expected, the ES model fits poorly: it essentially double counts effects in the high dose region where the dose-response curves flatten, a well known problem with this model.

**Figure 3.**
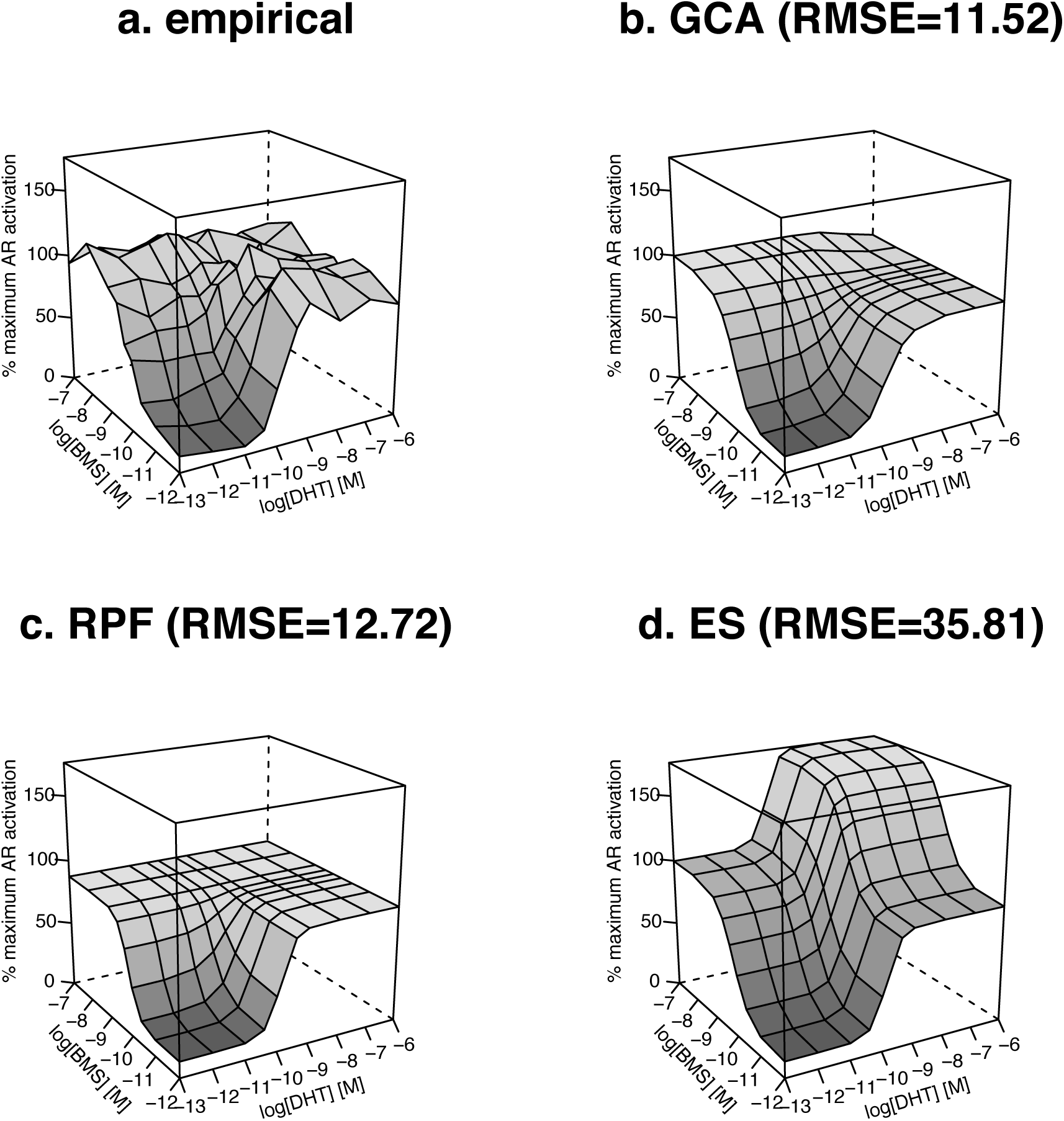
Response surface modeling of androgen receptor activation by a mixture of full agonists. Androgen receptor reporter data were generated from mixtures of DHT and BMS564929 as described in Figure 2A. Vertical axis: % Maximal AR Activation. Marginal curves correspond to individual dose-response curves shown in Figure 1 (shown as lowest concentration on log scale) except for RPF model. **(a)** Experimental joint dose-response surface. **(b)** GCA model based on parameters for PDM. **(c)** Relative Potency Factor model using DHT as the reference compound and Hill functions for the margins. **(d)** Effect Summation model using Hill functions for the margins. RMSE = root mean square error comparing fit of model to experimental data.

Figure 4 shows that for the full agonist/partial agonist mixture, GCA fits the experimental data quite well. The RPF model falls short due to the difference in efficacy between the two compounds. ES fits the data poorly as in Figure 3.

**Figure 4.**
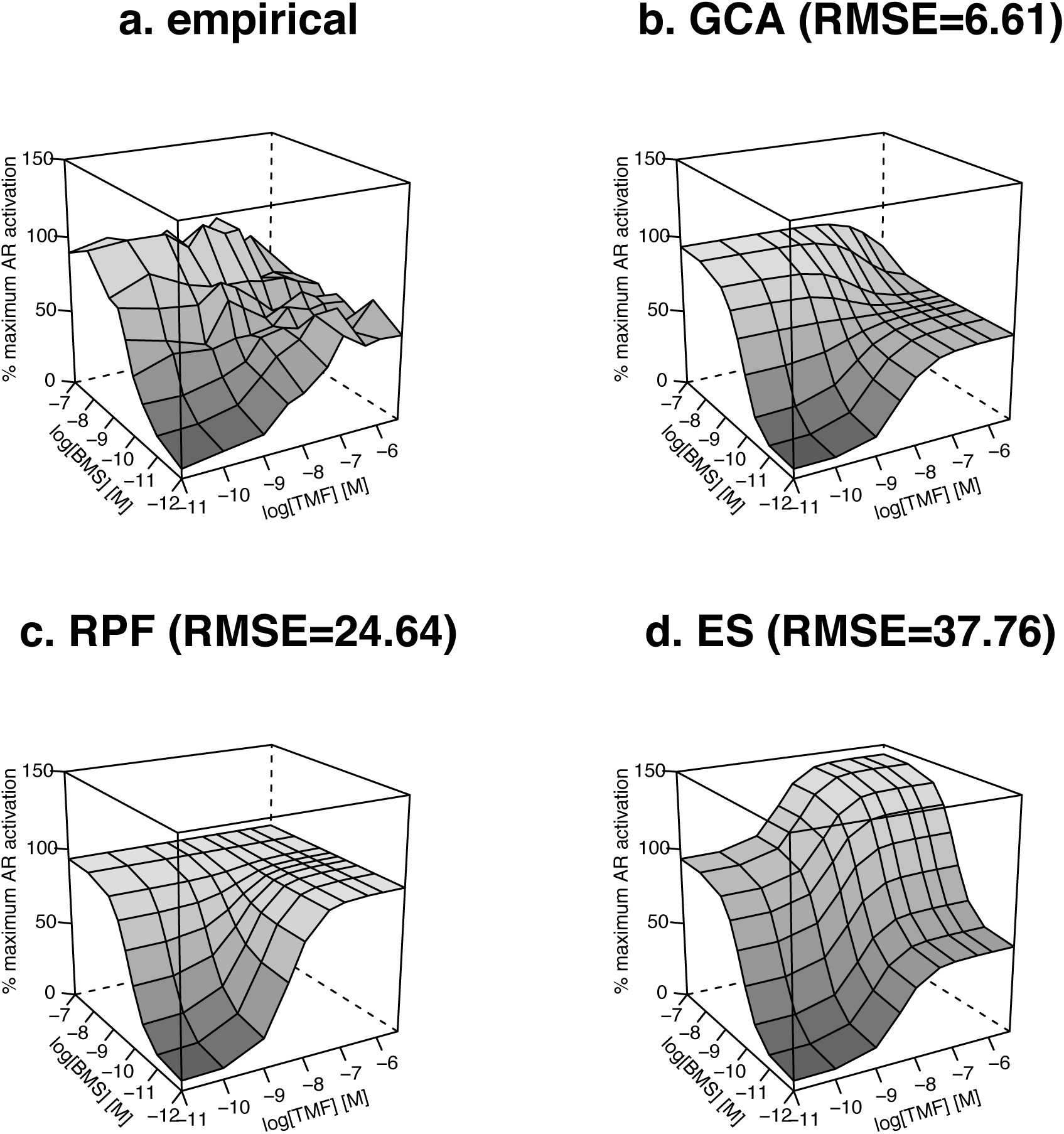
Response surface modeling of androgen receptor activation by a mixture of a full and a partial agonist. Androgen receptor reporter data were generated from mixtures of BMS564929 and TFM-4AS-1 as described in Figure 2B. Vertical axis: % Maximal AR Activation. Marginal curves correspond to individual dose-response curves shown in Figure 1 (shown as lowest concentration on log scale) except for RPF model. **(a)** Experimental joint dose-response surface. (b) GCA model based on parameters for PDM. **(c)** Relative Potency Factor model using BMS as the reference compound and Hill functions for the margins. **(d)** Effect Summation model using Hill functions for the margins. RMSE = root mean square error comparing fit of model to experimental data.

Next, we tested the applicability of GCA in modeling AR activation in the presence of a competitive antagonist. We began by establishing the dose-response of MDV3100; it fully antagonized AR activation by BMS564929 (Figure 5). MDV3100 is a competitive inhibitor of DHT binding to AR (Tran *et al.*, 2009). Therefore, we used the Schild method to estimate the equilibration dissociation constant for MDV3100 binding to AR in MDA-kB2 cells (Figure 6). We fit BMS564929 dose response curves generated in the presence of increasing concentrations of MDV3100 with a 4-parameter Hill model (Figure 6A) and a 2-parameter Hill model constrained to the mean slope and maxima (Figure 6B). Using Akaike’s information criteria (AICc, Figure 6C), we determined that the fit of the curves with the means generated a lower AIC; therefore, this model is statistically preferable. Using the EC_50_s in the dose response curves in Figure 6B, we calculated the dose ratios (DR) for each curve relative to the curve without MDV3100 and used these values to generate a Schild plot (Figure 6D). Since the 95% confidence interval of the slope (0.8663 to 1.038) included unity, the data were refit to a slope of 1, and the pK_b_ was estimated from the x intercept when y=0, yielding an estimate of 69 nM. In GCA, a competitive antagonist can be considered a partial agonist with zero efficacy. We generated three-dimensional response surfaces of AR activation by the full agonist and antagonist mixture, to compare the empirical dose response data with those predicted by different additive models. As shown in Figure 7, the GCA model predicts the experimental data much better than ES. The RPF model is not relevant for this case.

**Figure 5.**
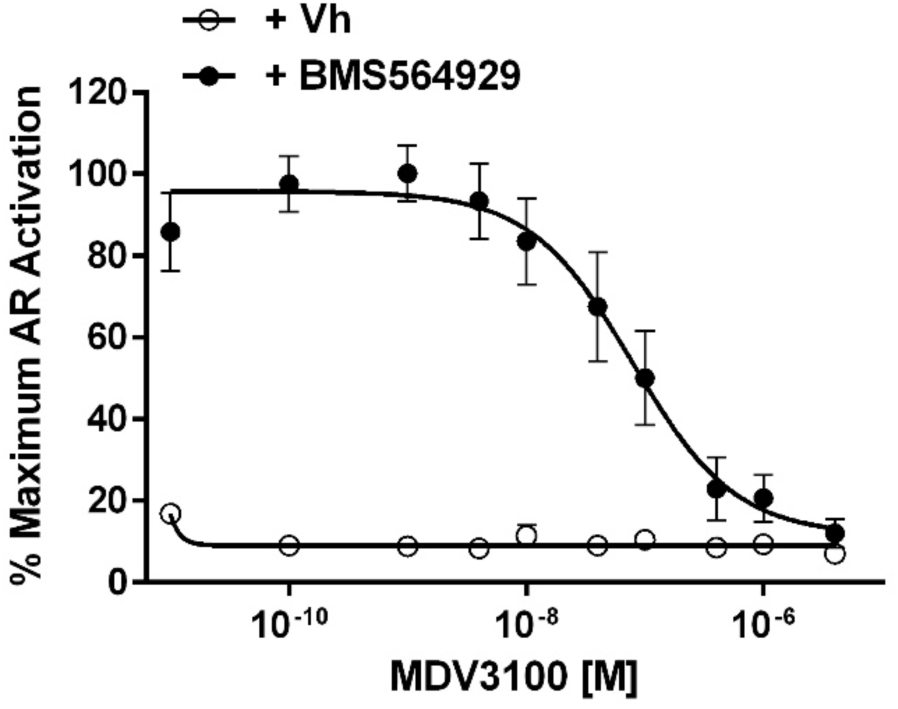
Dose response for androgen receptor activation by an antagonist in the absence and presence of a full agonist. MDA-kb2 cells were plated in steroid-free medium and dosed with Vh (0.5% DMSO), BMS564929 (1 × 10^−7^ M), and/or MDV3100 (1 × 10^−10^ M – 4 × 10^−6^ M). The cultures were then dosed with Vh (0.5% DMSO) or BMS564929 (1 × 10^−9^ M). Luminescence was measured after 24 hrs. Ligand-induced luminescence was normalized to the positive control-induced luminescence. Data from Vh-treated wells are reported as the lowest concentration in each plot. Data are presented as means ± SE. (N=6-10)

**Figure 6.**
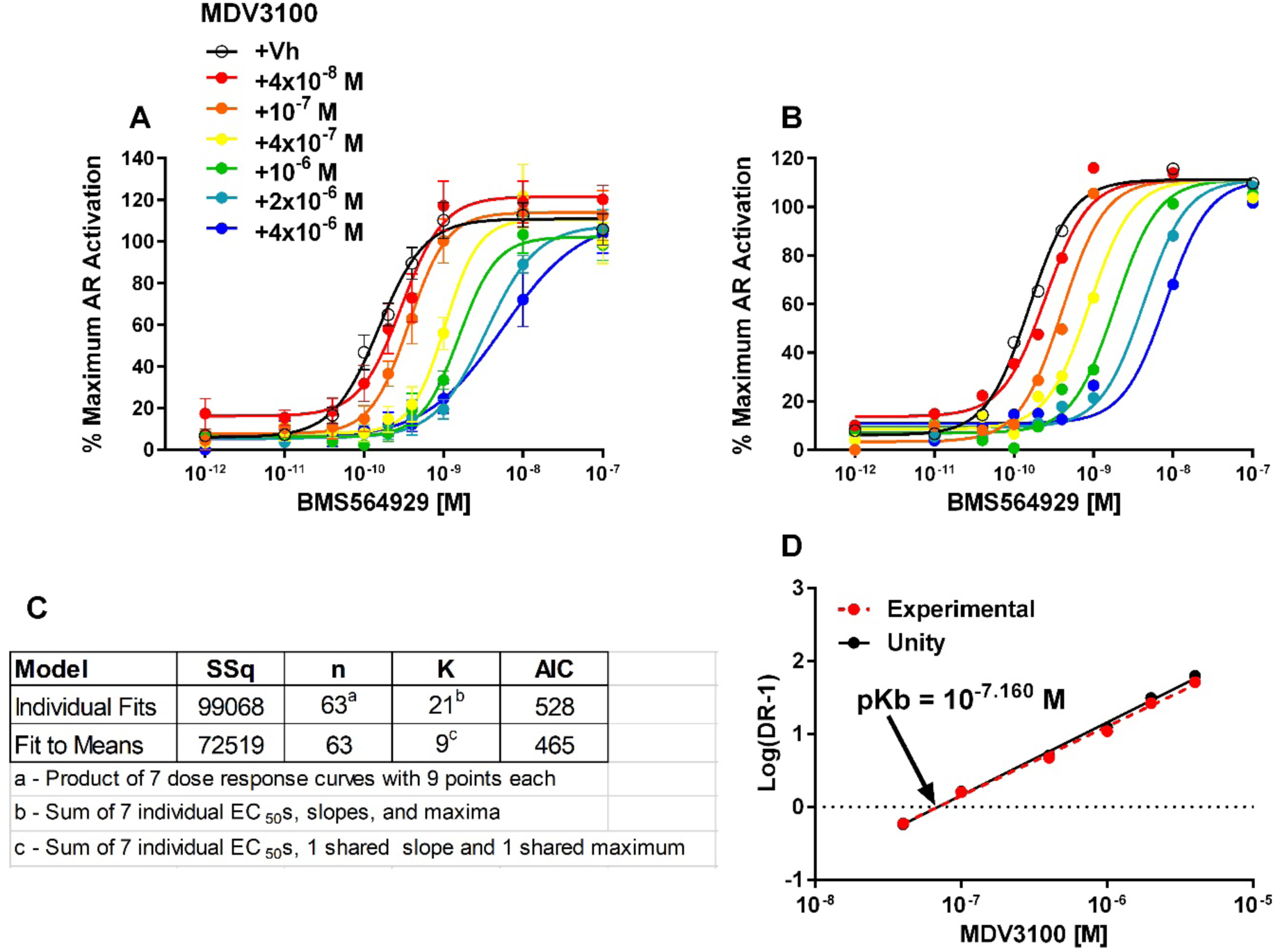
Schild analysis of MDV3100. MDA-kb2 cells were plated in steroid-free medium and dosed with Vh (0.5% DMSO), BMS564929 (1 × 10^−11^ M - 1 × 10^−7^ M), and/or MDV3100 (1 × 10^−10^ M – 2 × 10^−5^ M). Luminescence was measured after 24 hrs. Ligand-induced luminescence was normalized to the positive control-induced luminescence. Data from Vh-treated wells are reported as the lowest concentration in each plot. Data are presented as means ± SE. (N=6) **(A)** Individual dose response curves fitted with a 4 parameter Hill function. **(B)** Individual dose response curves fitted a 4 parameter Hill function constrained to the mean slope and maximum. **(C)** Akaike’s information criteria (AIC) for the individual and constrained fits. **(D)** Schild plot generated from the experimental data fit with the constrained Hill function and analyzed using linear regression. pKb is the estimate of the log of the dissociation constant of the competitive antagonist.

**Figure 7.**
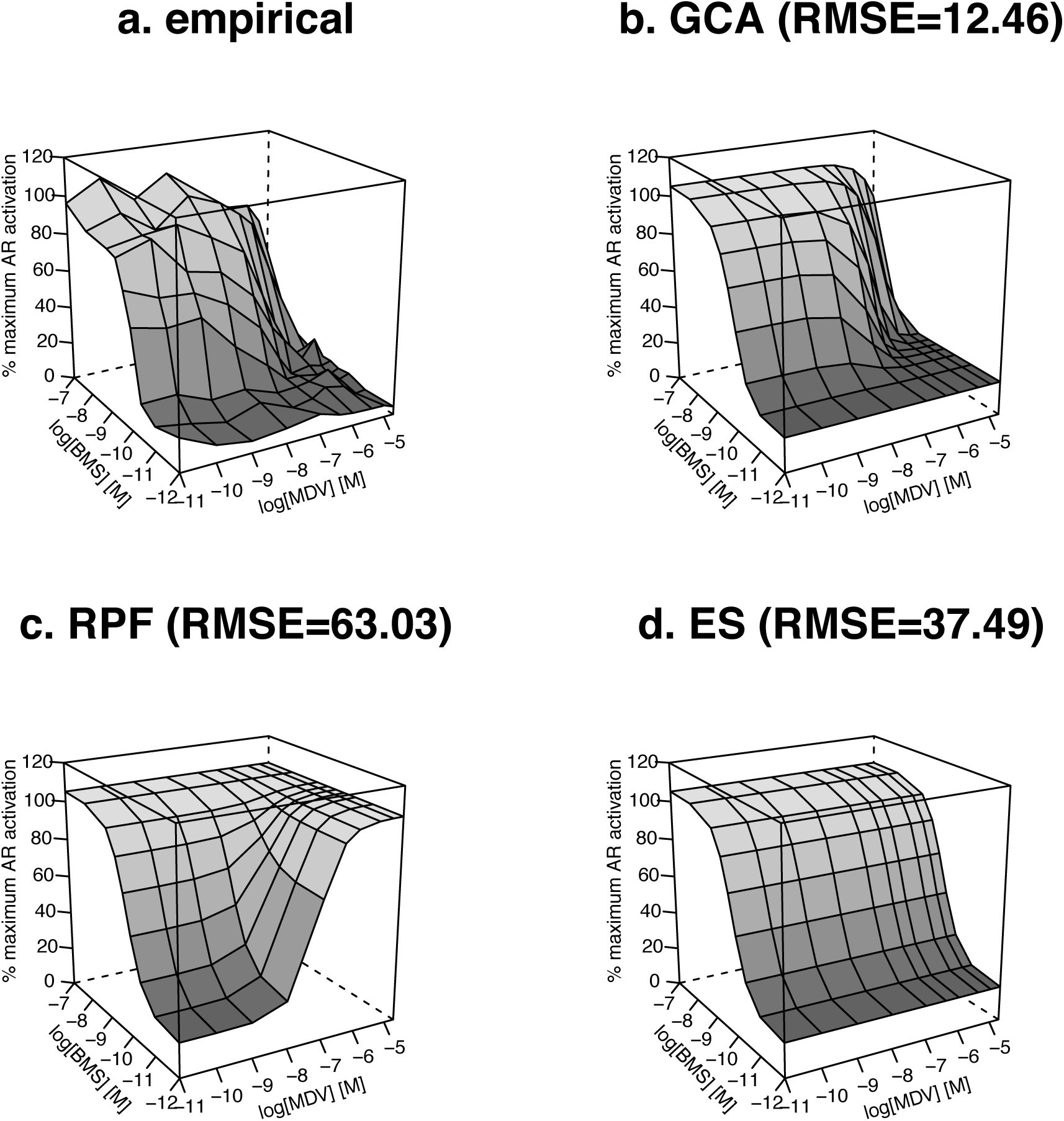
Androgen receptor activation by a mixture of a full agonist and a competitive antagonist. Androgen receptor reporter data were generated from mixtures of BMS564929 and MDV3100 as described in Figure 6. Vertical axis: % Maximal AR Activation. Marginal curves correspond to individual dose-response curves shown in Figure 1 for BMS and Figure 5 for MDV3100 (shown as lowest concentration on log scale) except for RPF model. **(a)** Experimental joint dose-response surface. **(b)** GCA model based on parameters for PDM and Schild analysis. **(c)** Relative Potency Factor model using BMS as the reference compound and Hill functions for the margins. **(d)** Effect Summation (ES).

## Discussion

Toxicologists typically study one compound at a time, but humans are exposed to a large number simultaneously. Furthermore, the National Institute of Environmental Health Science has made “unraveling the effects of environmental mixtures” a priority (Carlin *et al.*, 2013). The significance of NR-based mixture effects was underlined by research demonstrating that mixtures of environmental estrogens, each at its no observable effect level, elicited significant biological responses (Silva *et al.*, 2002). Here, we focused on the AR not only because of its physiological importance but also because it allowed us to test the efficacy of an expansion of our novel model of additivity, GCA, to a receptor system with two binding sites. We show that GCA can accommodate the pharmacodynamics of homodimers and accurately model AR activation by mixtures of full agonists, full and partial agonists and a full agonist and an antagonist.

We initially developed GCA in order to address a limitation of CA: it cannot model the effect of a mixture that contains ligands of differing efficacies at effect levels above that of the least efficacious mixture component (Silva *et al.*, 2002, Howard and Webster 2009). We showed that GCA can accurately model the activation of AhR and PPARγ by mixtures of ligands containing full and partial agonists (Howard, *et al*., 2010; Watt, *et al.*, 2016). While these receptors have distinct biological functions, they both can be modeled using a pharmacodynamic model for receptors with one binding site, a Hill function with a slope parameter of one. To expand the utility of GCA, we derived a pharmacodynamic model for homodimer receptors in which each monomer binds ligand and then dimerizes. The resulting function meets the mathematical requirements for application of GCA: the function is invertible yield real numbers. This property is not true for many commonly used empirical dose-response functions based on curve fitting, e.g., Hill functions with slope parameter greater than one (Webster and Schlezinger, 2019).

We tested the extension of GCA to modeling homodimers using empirical data generated with binary mixtures of ligands of the androgen receptor. The combination of two full agonists (DHT and BMS564929) generated mixture responses that were positively additive until the maximum activation level was reached; once the maximum was reached the mixture continued to fully activate the AR. GCA accurately modeled AR activation at all dose combinations; however, ES did not. As expected, ES overestimated the combined effect of the ligands at high concentrations. The RPF model, a special case of GCA and CA, performed nearly as well in this situation. The combination of a full and a partial agonist (BMS564929 and TFM-4AS-1) generated responses that were positively additive at low concentrations. At high combined concentrations, the activation of AR was reduced to the maximum AR activation induced by TFM-4AS-1 alone (i.e., ≈ 60% activation). This was predicted by GCA. Again, ES inadequately modeled the combined effect. The RPF model performed significantly worse than GCA as the assumption of equal efficacy is violated. These results indicate that GCA is a viable model in the case of two chemicals acting as either full or partial AR agonists.

Our ultimate goal is to use GCA to model AR activation by mixtures of natural ligands, pharmaceuticals, and environmental ligands, where many of the latter act as competitive antagonists (Luccio-Camelo and Prins, 2011). Thus, we next tested the ability of GCA to accommodate a mixture containing a competitive antagonist. Because the apparent IC_50_ of an antagonist depends upon the concentration of the agonist in the mixture, we used the Schild method to estimate the equilibrium constant for binding of a synthetic, competitive AR antagonist (Wyllie and Chen, 2007). The Schild method has been previously used to estimate the apparent binding affinity of a mixture of eight AR antagonists (Orton *et al.*, 2012). The linear shape of the Schild plot and the slope of ≈ 1 indicated that MDV3100 is a competitive AR antagonist. When the Schild method-derived equilibrium dissociation constant is incorporated into GCA, GCA accurately models the activation of AR by a mixture of a full agonist and an antagonist.

“Interaction” and “non-interaction” are ambiguous concepts in the absence of a definition of the latter. Our results suggest that ligands of the AR behave in a “non-interactive” manner using the definition of GCA that encompasses full and partial agonists as well as competitive antagonists. For example, we do not see evidence of cooperativity in the dimerization of AR when there are different ligands (Webster and Schlezinger 2019). We note that GCA may not apply where compounds act by different mechanisms (Webster, 2013), although there is some evidence that concentration addition performs better than the independent action model for the androgen system (e.g., NRC, 2008)

There are many chemicals in our indoor and outdoor environments, as well as in consumer products, that are AR ligands. They have the potential to disrupt both male and female reproductive health. We show that GCA can accommodate dose response functions for ligands (full agonists, partial agonists, and competitive antagonists) of homodimers, which significantly expands its utility. Future research will investigate the ability of GCA to model AR activation by complex (3+ components) environmental ligand mixtures and compare GCA with other approaches used to model AR (Birkhoj *et al.*, 2004; Blake *et al.*, 2010; Ermler et al., 2011; Ezechiáš and Cajthaml 2018; Kjeldsen *et al.*, 2013). Additionally, we are exploring the use of GCA to model other homodimer systems (de la Rosa et al, manuscript).

## Acknowledgments

This work was supported by the National Institute of Environmental Health Sciences grant R01 ES027813. The authors thank Mr. Nathan Burritt for his excellent technical assistance.

