## Supplemental Materials for "Predicting the activation of the androgen receptor by mixtures of ligands using Generalized Concentration Addition"

### Table of Contents

| | Page |
|---|---|
| Table S1. Parameter fits for individual compounds | 2 |
| Table S2. RMSE for 4 parameter Hill function and homodimer PDM fits | 2 |
| Figure S1. Toxicity analysis | 3 |

**Table S1. Parameter fits for individual compounds**

| | DHT | BMS | TMF |
|---|---|---|---|
| Hill function | | | |
| $a_0$ | 12.20 | 9.888 | 9.884 |
| span | 75.26 | 87.47 | 49.85 |
| $EC_{50}$ | 1.261 E-10 | 1.400 E-10 | 3.968 E-9 |
| p | 2.046 | 1.295 | 1.315 |
| PDM* | | | |
| α | 8.567 | 24.22 | 3.048 |
| K | 3.921 E-10 | 1.208 E-9 | 4.741 E-9 |

* The PDM fit used the a0 from the Hill fit and a common λ=95

**Table S2. RMSE for 4 parameter Hill function and homodimer PDM fits**

| | DHT | BMS | TMF |
|---|---|---|---|
| Hill | 16.85 | 15.56 | 14.52 |
| PDM | 17.19 | 15.57 | 14.57 |

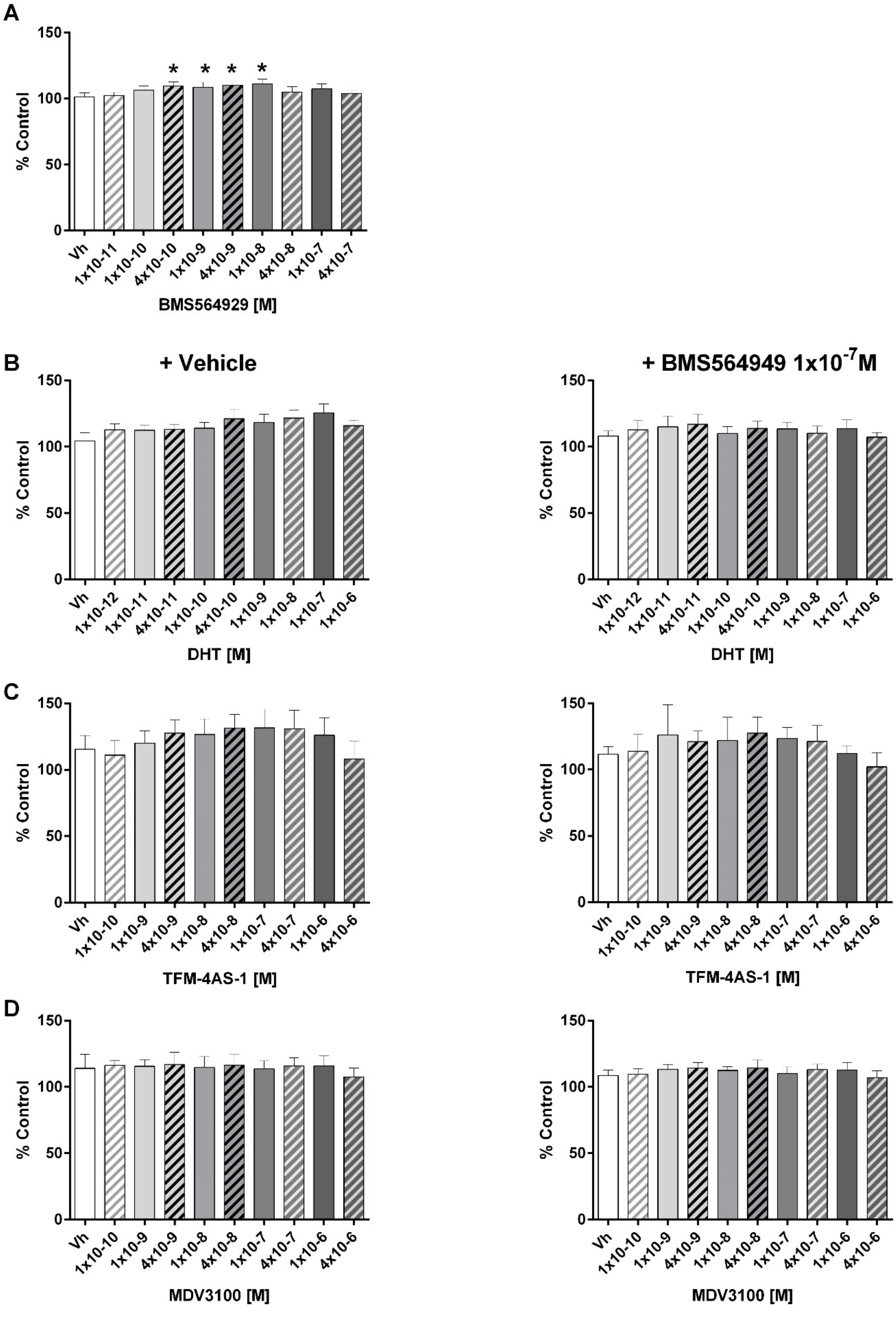
A
B
C
D
+ Vehicle
+ BMS564949 1x10-7M
% Control
0
50
100
150
*
Vh
1x10-11
1x10-10
4x10-10
1x10-9
4x10-9
1x10-8
4x10-8
1x10-7
4x10-7
BMS564929 [M]
1x10-12
4x10-11
1x10-6
DHT [M]
4x10-7
4x10-6
TFM-4AS-1 [M]
MDV3100 [M]

**Figure S1. Toxicity analysis.** MDA-kb2 cells were plated in steroid-free medium, treated with Vh (0.5% DMSO), **(A)** BMS564929 (1 x $10^{-11}$ M – 4 x $10^{-7}$ M), **(B)** DHT (1 x $10^{-12}$ M – 1 x $10^{-6}$ M), **(C)** TFM-4AS-1 (1 x $10^{-10}$ M – 4 x $10^{-6}$ M), or **(D)** MDV3100 (1 x $10^{-10}$ M – 4 x $10^{-6}$ M), co-treated with Vh or BMS564929 (1 x $10^{-7}$ M). Toxicity was analyzed after 24 hrs. Absorbance in experimental wells was normalized to absorbance in wells treated with $1x10^{-8}$ M DHT (Control). Data are presented as means ± SE of at least 4 independent experiments. * Significantly different from Vh ($p < 0.05$, ANOVA, Dunnett's).
